# A computational strategy to uncover fusion genes in prostate cancer cell lines

**DOI:** 10.1101/2025.05.12.653554

**Authors:** Rebecca A. Morgan, Gary Hardiman

**Author notes:** Correspondence should be addressed to Gary Hardiman.

## Abstract

Fusion genes, chimeric transcripts formed by the merging of two distinct genes due to chromosomal structural changes (e.g., inversions or trans/cis-splicing), are established cancer drivers. Advances in genomic technologies, particularly RNA sequencing and improved fusion gene prediction algorithms, have significantly expanded our understanding of fusion genes in cancer. This chapter explores computational methods for identifying fusion genes through RNA sequencing data, using the *TMPRSS2::ERG* fusion in prostate cancer cell lines as a case study, and includes analysis of both fusion-positive and fusion-negative cell lines. To achieve high-confidence detection, three open-source fusion prediction tools, STAR-Fusion, FusionCatcher, and JAFFA are investigated. These tools were selected for their accessibility, active maintenance, and strong performance in benchmarking studies. Their sensitivity and accuracy in detecting *TMPRSS2::ERG* is systematically evaluated and validated, ensuring robust and high-resolution detection of fusion events.

## 1. Introduction

Fusion genes are formed by the combination of two genes due to chromosome breakage and re-splicing at the genomic level ***(1)***. These fusion genes are transcribed and translated into a single unit, producing novel proteins with unique properties that differ from those encoded by the original genes ***(2, 3)***. Identified as a driving force behind malignancy development, fusion genes play a critical role in cancer progression ***(4)***. It has been estimated that fusion genes account for approximately 20% of cancer morbidity ***(5, 6)***, however fusion gene incidence rates considerably differ across cancer types with many recurrent fusion genes identified ***(5)***, examples include *BCR::ABL1* found to occur in more than 95% of chronic myeloid leukaemia patients ***(1)***, *EML4::ALK* estimated to occur in 2-9% of lung adenocarcinomas ***(7)***and *PAX3::FOX01* found to occur in 60% of alveolar rhabdomyosarcoma cases ***(8)***. The fusion between the prostate-specific androgen-responsive gene Transmembrane protease, serine 2 (*TMPRSS2*) and the ETS-related gene (*ERG*) has been widely investigated in prostate cancer (PC) [15,16]. *TMPRSS2::ERG* in prostate cancer primarily drives ERG overexpression, which activates critical biological pathways such as androgen receptor signaling, cell cycle regulation, DNA damage repair, and epithelial-to-mesenchymal transition. These pathways and their associated proteins collectively contribute to the progression and aggressiveness of prostate cancer [16]. High-Throughput Sequencing (HTS) has revolutionized the detection of fusion genes, enabling rapid identification at both the transcript and genomic levels. As a result, fusion prediction software has rapidly evolved, driven by advances in computational biology and HTS. In this chapter, we evaluate and compare three open-source tools designed to detect *TMPRSS2::ERG* fusions in RNAseq data from prostate cancer cell lines. We outline the process from the generation of ‘FASTQ’ files by the sequencing instrument to the final list of predicted fusion genes for each cell line. All fusion prediction tools require working in a Linux environment. A basic knowledge of the command line environment, file structures and coding skills are assumed.

## 2. Materials and Methods

### 2.1. Computing environment

Computational fusion prediction analyses are intricate, multi-step processes that require integrating multiple software applications. Many fusion prediction programs are optimised for high-performance computing environments. For optimal performance, fusion prediction software requires a well-configured and powerful desktop. A system with an Intel Core i7 processor and storage configured in RAID 0 with two parallel 1 TB hard drives is recommended. Additionally, most fusion prediction programs need a minimum of 24GB of RAM, though 18GB may suffice for very small FASTQ files.

### 2.1. Software tools

Cutadapt (https://cutadapt.readthedocs.io),

Python. Windows/Linux/MAC OS X.

STAR-Fusion (https://github.com/STAR-Fusion/STAR-Fusion/wiki), Linux/MAC OS X.

FusionCatcher (https://github.com/ndaniel/fusioncatcher), Linux.

JAFFA (https://github.com/Oshlack/JAFFA), Linux.

### 2.2. Fusion gene prediction tools

Fusion prediction tools have emerged as an area of rapid growth with the expansion of HTS. RNAseq is the preferred method for fusion gene detection due to its lower cost, faster computational processing, and reduced memory requirements ***(9)***. Thus, most fusion prediction software is based on the analysis of RNAseq data. Despite the many fusion gene prediction tools developed, many have become obsolete or have not been updated. **Table 1** provides an overview of fusion gene prediction tools available, including their year of release, input data, read format, reference, URL link and status.

**Table 1.** Overview of RNAseq based fusion prediction tools.

| *Tool* | *Release year* | *Input data* | *Input format* | *Reference* | *URL* | *Status* |
| --- | --- | --- | --- | --- | --- | --- |
| **STAR-Fusion** | 2017 | RNAseq | Paired end | (10) | https://github.com/STAR-Fusion/STAR-Fusion | Active |
| **Fusion Catcher** | 2012 | RNAseq | Paired end | (11) | https://github.com/ndaniel/fusioncatcher | Active |
| **JAFFA** | 2015 | RNAseq | Single and paired end | (12) | https://github.com/Oshlack/JAFFA | Active |
| **Chimera Scan** | 2011 | RNAseq | Paired end | (13) | https://code.google.com/archive/p/chimerascan/ | Dormant |
| **EricScript** | 2012 | RNAseq | Paired end | (14) | https://github.com/databio/ericscript | Dormant |
| **SOAPFuse** | 2013 | RNAseq | Paired end | (15) | http://soap.genomics.org.cn/soapfuse.html | Dormant |
| **TopHat-Fusion** | 2011 | RNAseq | Single and paired end | (16) | http://tophat-fusion.sourceforge.net/ | Dormant |
| **Fusion Map** | 2011 | RNAseq and gDNA-seq | Single and paired end | (17) | http://www.omicsoft.com/fusionmap | Dormant |
| **Fusion Finder** | 2012 |  | Single and paired end | (18) | http://bioinformatics.childhealthresearch.org.au/software/fusionfinder/ | Dormant |
| **FuSeq** | 2018 | RNAseq | Paired end | (19) | https://github.com/nghiavtr/FuSeq | Active |
| **Fusion Scan** | 2019 | RNAseq | Single and paired end | (20) | http://fusionscan.ewha.ac.kr/ | Dormant |
| **Break Fusion** | 2012 | RNAseq | Paired end | (21) | https://bioinformatics.mdanderson.org/public-software/breakfusion/ | Active |
| **deFuse** | 2011 | RNAseq | Paired end | (22) | https://github.com/amcpherson/defuse | Active |

Based on each tool’s development date, acceptance within the research community (measured by citation frequency), current maintenance status (active web links), and ease of installation, STAR-Fusion, FusionCatcher, and JAFFA were chosen as the most suitable tools for the evaluation in this chapter.

#### 2.2.1. STAR-Fusion

STAR-Fusion, released in 2017, is a fusion detection tool that utilises a Python script for installation ***(10)***. It employs the STAR aligner and uses the transcriptome as a reference. The tool supports both single-end and paired-end read formats and is classified under read mapping, with RNAseq as its input data. One of its key advantages is its fast fusion discovery time, leveraging the transcriptome for efficient detection. However, a notable limitation is that it relies solely on the STAR aligner, which may impact its versatility compared to multi-aligner approaches.

#### 2.2.2. FusionCatcher

FusionCatcher, released in 2012, is a fusion detection tool that installs via a Python script and utilises multiple alignment tools, including Bowtie, Bowtie2, Liftover, STAR, Velvet, Fatotwobit, SAMtools, Seqtk, Numpy, Biopython, Picard, and Parallel ***(11)***. Unlike some tools that rely solely on a transcriptome reference, FusionCatcher uses both the genome and transcriptome for improved detection accuracy. It supports paired end read formats and falls under the read mapping classification, processing RNAseq input data. One of its key advantages is the inclusion of assembly software, which enhances fusion detection, along with its high recall rates. However, a notable drawback is the presence of hard-set false and positive fusions, which may impact result precision.

#### 2.2.3. JAFFA

JAFFA, released in 2015, is a fusion detection tool that installs via a Java script and utilises multiple alignment tools, including Bpipe, Velvet, Oases, SAMtools, Bowtie2, BLAT, Dedupe, and Reformat ***(12)***. It relies on the transcriptome as its reference and supports both single-end and paired-end read formats. Unlike some tools that focus solely on read mapping, JAFFA combines both read mapping and assembly approaches, processing RNAseq input data. Its key advantages include the incorporation of assembly software, the ability to run multiple samples in parallel, and support for both paired and single-end data. However, these capabilities come at the cost of higher computational time and memory usage. Additionally, JAFFA is not highly sensitive to intronic fusions, which may limit its detection scope.

### 2.3. Cell lines

RNA sequencing data from prostate cancer cell lines, known to be either positive or negative for the *TMPRSS2::ERG* fusion, were analysed to assess the presence of this fusion. The sequence data were obtained from the NCBI Sequence Read Archive (accession code PRJNA523380)

https://www.ncbi.nlm.nih.gov/bioproject/?term=PRJNA523380.

The data can also be obtained from

https://www.ebi.ac.uk/ena/browser/view/SRR8616157

All RNAseq data for the cell lines are available in paired end format. A summary of the cell lines examined is presented in Table 2. To evaluate detection accuracy, we determined whether each tool identified the fusion in accordance with the known *TMPRSS2::ERG* status of each cell line. The analysed cell lines included VCaP (+), HCIH-660 (+), PC3 (−), LNCaP (−), DU145 (−), and 22RV1 (−), along with PrECLH and MDA-PCa-2B, whose status is unknown. This approach provided both positive and negative controls for *TMPRSS2::ERG* detection, enabling an assessment of the fusion prediction tools’ accuracy in the context of PC. HCIH-660 and VCaP served as positive controls, while PC3, LNCaP, DU145, and 22RV1 were used as negative controls.

**Table 2.**
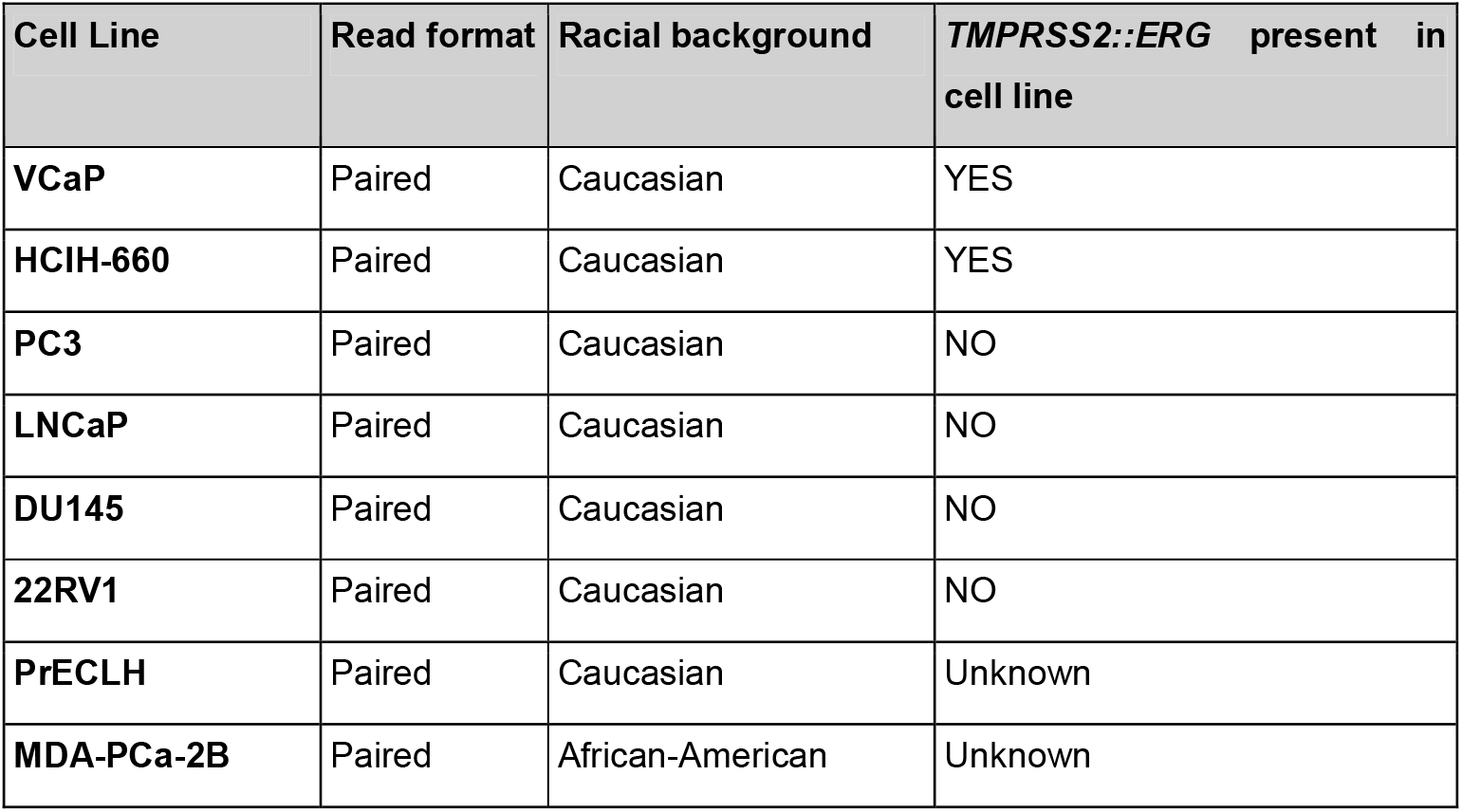
Summary of prostate cancer model cell lines.

| Cell Line | Read format | Racial background | *TMPRSS2::ERG* present in cell line |
| --- | --- | --- | --- |
| VCaP | Paired | Caucasian | YES |
| HClH-660 | Paired | Caucasian | YES |
| PC3 | Paired | Caucasian | NO |
| LNCaP | Paired | Caucasian | NO |
| DU145 | Paired | Caucasian | NO |
| 22RV1 | Paired | Caucasian | NO |
| PrECLH | Paired | Caucasian | Unknown |
| MDA-PCa-2B | Paired | African-American | Unknown |

## 3. Methods

Figure 1 illustrates a schematic representation of the data analysis pipeline. For this exercise, FASTQ files can be downloaded via the European Nucleotide Archive (https://www.ebi.ac.uk/ena/browser/view/SRR8616157 using the ‘wget’ command. Readers are recommended to check the quality of the FASTQ files prior to beginning fusion analysis. This can be performed using the FastQC software available from https://www.bioinformatics.babraham.ac.uk/projects/fastqc/. All samples are paired end. Paired-end sequencing samples follow a standard naming convention, with SRR8615300_1.fastq.gz representing read one (R1) and SRR8615300_2.fastq.gz representing read two (R2) etc.

**Figure 1.**
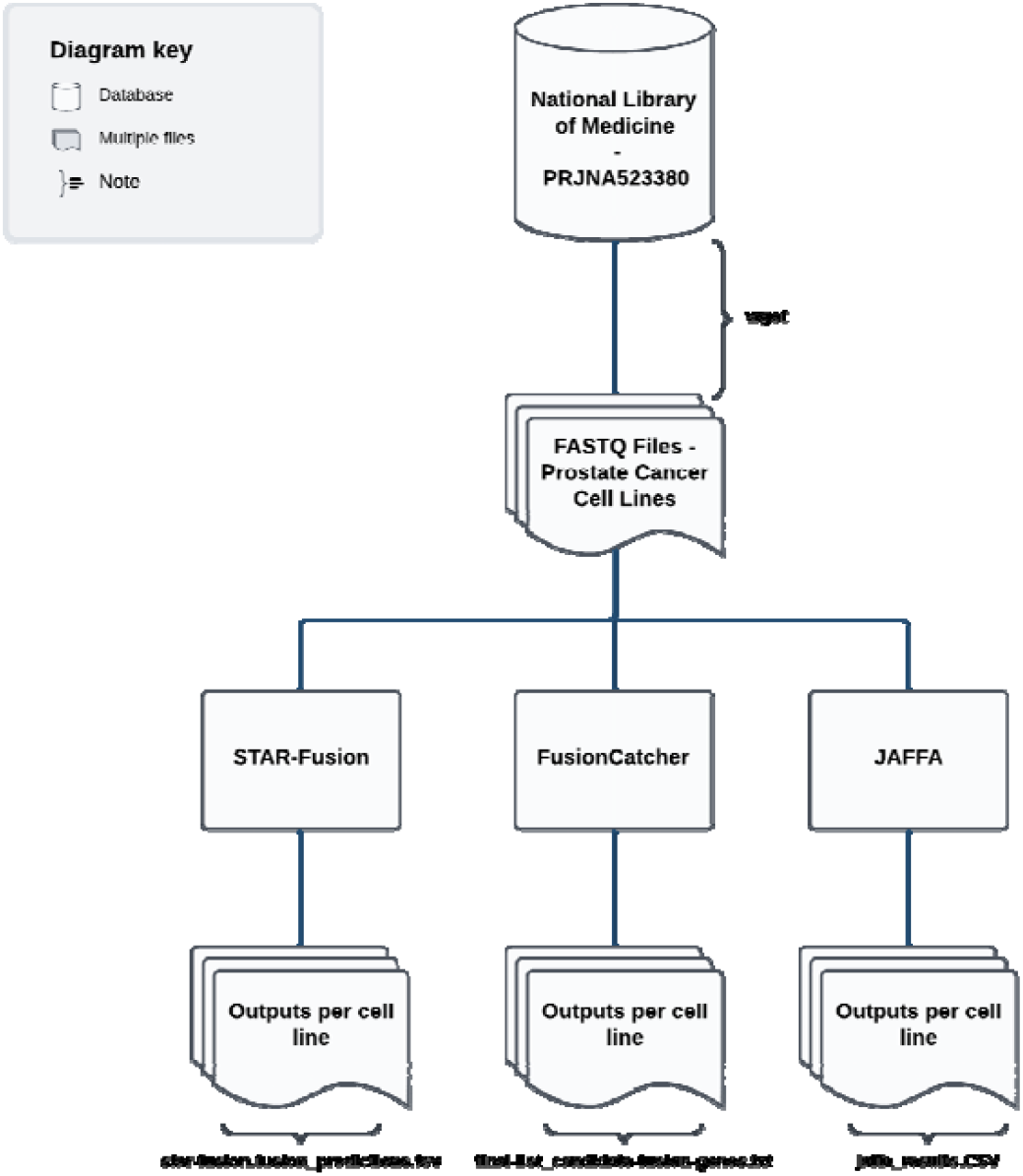
Flowchart summarising the data analysis process, from the collection of FASTQ files to fusion event predictions for each tool.

### 3.1. STAR-Fusion

#### 3.1.1. Code

The installation guide for STAR-Fusion is available on https://github.com/STAR-Fusion/STAR-Fusion/wiki/installing-star-fusion. STAR-Fusion requires use of the STAR alignment tool to predict fusion events. Users are required to choose a reference genome. For this chapter, GRCh38 was selected as the reference genome. Unlike FusionCatcher and JAFFA, STAR-Fusion does not automatically identify all paired FASTQ files from a specified directory. To streamline the processing of these samples, a Bash script can be used to automate fusion gene detection by identifying and analysing all paired FASTQ files in the current directory. The script, shown in Code Snippet A, contains a for loop that iterates through _1.fastq.gz files, extracts the base name, identifies the corresponding _2.fastq.gz file, and creates a results directory for each sample. It then executes STAR-Fusion using the identified read pairs, specifying the genome library directory and an output location. Alternatively, users can analyse paired samples individually, as demonstrated in Snippet B for the cell line DU145 (SRR8615300).

**A.**

~~~
#!/bin/bash
# Activate STAR-Fusion environment
conda activate star-fusion
# Create output directory if it doesn’t exist
mkdir -p /mnt/rebecca/star-fusion_cell_line_exercise_results
# Loop through all _1.fastq.gz files
for i in *_1.fastq.gz; do
  # Extract the base file name (without _1.fastq.gz)
  fname=$(basename --“ $i”)
  fnamenoEnd=${fname%_1.fastq.gz}
  R2=${fnamenoEnd}_2.fastq.gz # Define paired read file
# Create an output directory for this sample
mkdir -p /mnt/rebecca/star-fusion_cell_line_exercise_results/” $fnamenoEnd”
# Run STAR-Fusion
STAR-Fusion --genome_lib_dir
  /mnt/hdd/rebecca/_star_fusion_ctat/GRCh38_gencode.plug-n-
play/ctat_genome_lib_build_dir/ \
    --left_fq “ $i” \
    --right_fq “ $R2” \
    --output_dir /mnt/rebecca/star-fusion_cell_line_exercise_results /” $fnamenoEnd”
done
~~~

**B.**

~~~
#!/bin/bash
conda activate star-fusion
# Create output directory if it doesn’t exist
mkdir -p /mnt/rebecca/star-fusion_cell_line_exercise_results
STAR-Fusion --/mnt/hdd/rebecca/_star_fusion_ctat/GRCh38_gencode.plug-n-
play/ctat_genome_lib_build_dir \
    --left_fq SRR8615300_1.fastq.gz \
    --right_fq SRR8615300_2.fast.gz \
    --output_dir /mnt/rebecca/star-fusion_cell_line_exercise_results
~~~

#### 3.1.2. STAR-Fusion results

STAR-Fusion generates multiple output files for each cell line, but the key file for this exercise is star-fusion.fusion_predictions.tsv, which contains a list of predicted fusion events. Readers are advised to open this file in Microsoft Excel and refer to the #FusionName column for a list of predicted fusion genes. https://github.com/STAR-Fusion/STAR-Fusion/wiki#output-from-star-fusion provides a full description of the column headers in this output file. Figure 2 shows an example of the star-fusion.fusion_predictions.tsv file for the VCap cell line (SRR8618305) opened in Excel.

**Figure 2.**
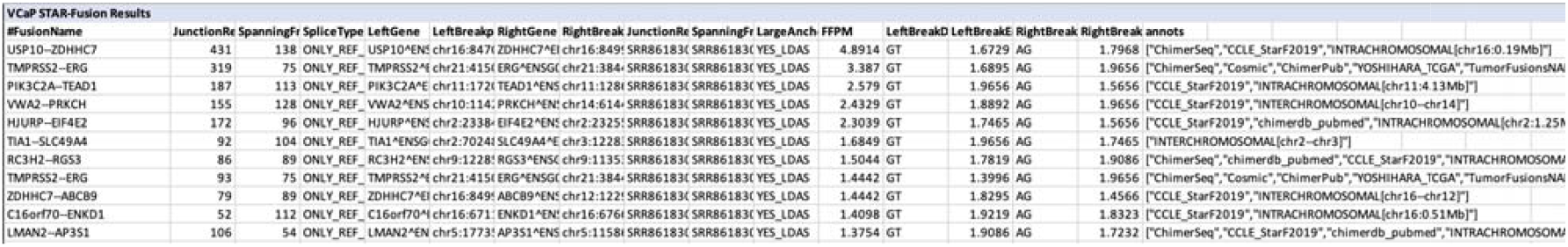
STAR-Fusion results (star-fusion.fusion_predictions.tsv).

### 3.2. FusionCatcher

#### 3.2.1. Code

The installation guide for FusionCatcher is available from https://github.com/ndaniel/fusioncatcher/blob/master/doc/manual.md#4---installation-and-usage-examples. This documentation also includes information on how to download a chosen reference genome. For this exercise, GRCh38 was selected. Unlike STAR-Fusion, FusionCatcher automatically detects paired reads when provided with a folder containing all read pairs. To ensure proper detection, sample names must be synchronised and correctly labelled as R1 or R2. In this case, the cell line samples are already formatted correctly and require no modifications. Code snippet C provides a script which executes FusionCatcher using a predefined reference genome directory and the specified output location.

**C.**

~~~
#!/bin/bash
# Add conda environment to path
export PATH=“ $PATH:/usr/local/bin/conda/envs/integrate_soap/bin/”
# Run FusionCatcher
python /usr/local/bin/conda/envs/integrate_soap/bin/fusioncatcher.py \
-d /mnt/hdd/rebecca/FusionCatcher_genome_build/current/ \
-i /mnt/hdd/exercise/cell_lines_PC/ \
-o                /mnt/hdd/exercise/cell_lines_PC/results/-o
/mnt/hdd/exercise/cell_lines_PC/FusionCatcher_results/
~~~

#### 3.2.2. FusionCatcher results

FusionCatcher generates multiple output files for each cell line, but the key file for this exercise is final-list_candidate_fusion_genes.txt, which contains a list of predicted fusion events. Readers are advised to open this file in Microsoft Excel and refer to columns Gene_1_symbol(5end_fusion_partner) and Gene_2_symbol(3end_fusion_partner) for a list of predicted fusion genes. Figure 3 shows an example of the final-list_candidate_fusion_genes.txt file for the VCap cell line (SRR8618305) opened in Excel. https://github.com/ndaniel/fusioncatcher/blob/master/doc/manual.md#62---output-data-output-data provides a full description of the column headers in this output file.

**Figure 3.**
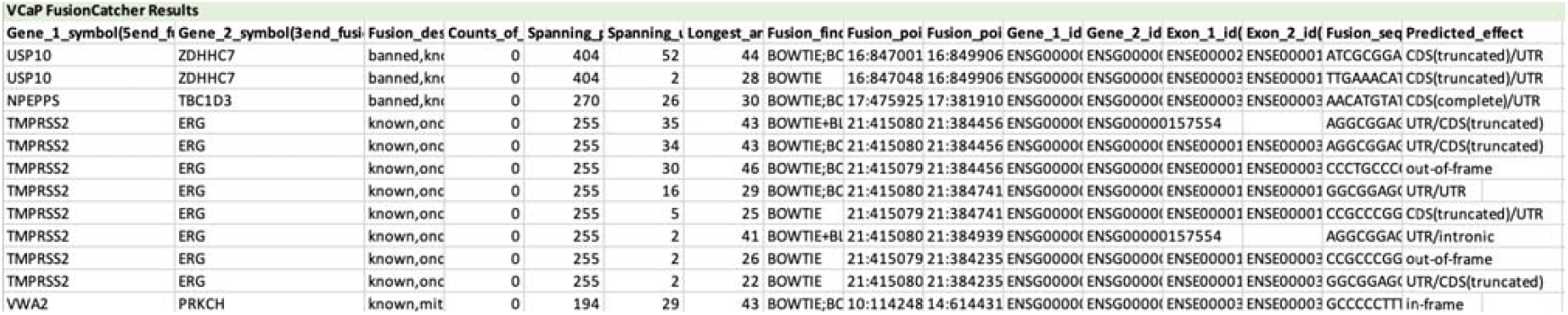
FusionCatcher results (final-list_candidate_fusion_genes.txt).

### 3.3. JAFFA

#### 3.3.1. Code

This Bash script executes the JAFFA Direct pipeline using Bpipe, a bioinformatic workflow system that consists of a series of processing steps ***(23)***. It begins by calling the Bpipe platform from its installation directory to run the JAFFA Direct pipeline. JAFFA offers users three modes: Assembly, Direct and Hybrid. More information on the difference between these three modes are available in the 2015 paper by Davidson et al ***(12)***. For this, exercise, Direct mode was utilised. The JAFFA Direct pipeline script (JAFFA_direct.groovy) is provided as an argument, along with all gzip-compressed FASTQ files located in the /mnt/hdd/exercise/cell_lines_PC/ directory. Bpipe orchestrates the execution of JAFFA, ensuring proper processing of input files while leveraging parallel computing to speed up the analysis. Code snippet D provides a script for running JAFFA in direct mode with 10 processing cores.

**D.**

~~~
#!/bin/bash
/mnt/hdd/jaffa/JAFFA-version-1.09/tools/bpipe-0.9.9.2/bin/bpipe run -n 10 \
/mnt/hdd/jaffa/JAFFA-version-1.09/JAFFA_direct.groovy \
/mnt/hdd/exercise/cell_lines_PC/*.gz
~~~

#### 3.3.2. JAFFA results

JAFFA generates multiple output files for each cell line, but the key file for this exercise is jaffa_results.CSV, which contains a list of predicted fusion events. Readers are advised to open this file in Microsoft Excel and refer to column fusion_genes for a list of predicted fusion genes. JAFFA provides users with the option to filter fusions based on classification (high, medium or low confidence). A description of these different classification types is provided in the paper by Davidson and colleagues***(12)***. For this exercise, only high confidence fusion events were selected. Figure 4 shows an example of the jaffa_results.CSV file for the VCap cell line (SRR8618305) opened in Excel and filtered by the high confidence classification. https://github.com/Oshlack/JAFFA/wiki/OutputDescription#jaffa_resultscsv provides a full description of the column headers in this output file.

**Figure 4.**
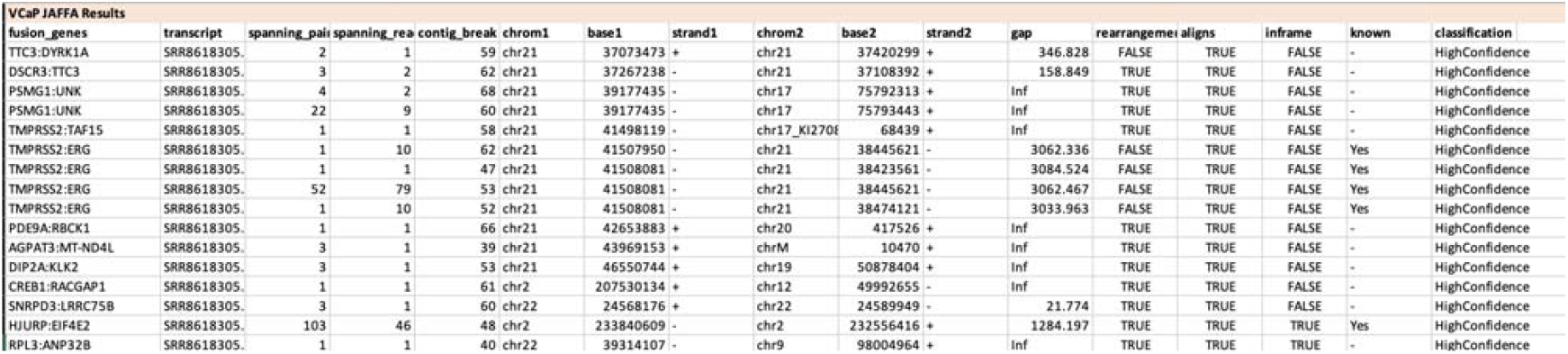
JAFFA results (jaffa_results.CSV).

## 4. Results

### 4.1. Benchmarking

*TMPRSS2::ERG* was identified in the positive control cell lines (VCaP and HCIH-660) using STAR-Fusion, FusionCatcher, and JAFFA. JAFFA detected this fusion with high confidence in both VCaP and HCIH-660. None of the tools predicted *TMPRSS2::ERG* in the negative control cell lines. Overall results indicate STAR-Fusion, FusionCatcher and JAFFA provide reliable methods for predicting fusion events. Table 3 presents a summary of the benchmarking of STAR-Fusion, FusionCatcher and JAFFA. Prostate cancer is notably prevalent among men of African descent and those living a Western lifestyle, indicating a mix of genetic, environmental, and socioeconomic risk factors. Advances in RNA sequencing and fusion gene detection algorithms have enhanced our understanding of cancer, particularly through identifying fusion genes, which are significant drivers of the disease. These tools can be expanded beyond the benchmarking provided to investigate molecular differences in fusion genes between African American and European American men, revealing unique gene fusions***(24)***.

**Table 3.** Cell line results by tool.

| Cell Line | *TMPRSS2::ERG*<br>present in cell line | Predicted by STAR-Fusion | Predicted by FusionCatcher | Predicted by JAFFA |
| --- | --- | --- | --- | --- |
| VCaP | YES | YES | YES | YES |
| HClH-660 | YES | YES | YES | YES |
| **PC3** | NO | NO | NO | NO |
| **LNCaP** | NO | NO | NO | NO |
| **DU145** | NO | NO | NO | NO |
| **22RV1** | NO | NO | NO | NO |
| **PrECLH** | Unknown | NO | NO | NO |
| **MDA-PCa-2B** | Unknown | NO | NO | NO |

## Conclusions

This chapter has demonstrated the efficacy of contemporary computational tools in detecting fusion genes, pivotal cancer drivers formed by chromosomal anomalies such as inversions or complex splicing events. Through a focused study on the *TMPRSS2::ERG* fusion in prostate cancer cell lines, we assessed the performance of three open-source fusion prediction tools: STAR-Fusion, FusionCatcher, and JAFFA. Our findings confirm that these tools effectively identify *TMPRSS2::ERG* fusions in known positive control cell lines, VCaP and HCIH-660, with JAFFA showing particularly high confidence in its predictions. Importantly, all tools consistently showed no false positives in the negative control cell lines, underscoring their reliability and precision in fusion detection. This systematic evaluation not only enhances our understanding of the capabilities and limitations of these computational approaches but also reinforces the role of advanced RNA sequencing technologies in advancing the molecular characterization of cancer, potentially leading to improved diagnostic and therapeutic strategies.

